# Determine Interaction Affinity Changes of the SUMO E1 Activating Enzymes During SUMO Activation using Quantitative FRET technology

**DOI:** 10.1101/2025.07.30.667747

**Authors:** Ling Jiang, Yiran Tao, Xin Wen, Jiayu Liao

## Abstract

The heterodimeric E1 complex, Aos1-Uba2, catalyzes the first adenylation activation of the SUMO1 peptide in the SUMOylation cascade. The reaction affinity and dynamics of the Aos1-Uba2 heterodimer during the first step activation have yet to be determined. The Kd for the Aos1-Uba2 interaction provides a unique perspective for the activation step of the ubiquitin-like protein conjugation cascade. Here, we report for the first time the determination of the Aos1 and Uba2 interaction dissociation constant (Kd) and kinetics using the qFRET assay. We also used the SPR method to verify the interaction Kd between Aos1 and Uba2. We also determined the kinetics changes of Aos1-Uba2 when SUMOs and ATP were added to the reaction in real time. The results showed that forming a thioester bond between SUMO1 and Uba2 increases the FRET signal, indicating that the E1 heterodimer is more stable and bound to each other in SUMO and ATP. These results suggest that the qFRET method can be used to determine protein interaction affinity changes and track real-time changes in protein conformation and dynamics changes during biochemical reactions.

## Introduction

The ubiquitin-like (UBL) protein modification, an important post-translational pro-tein modification for almost one-third of human proteins, plays critical roles in diverse physiological processes, such as transcriptional regulation, signal transduction, cell sur-vival and death, DNA damage responses, and spindle organization1,2. The small ubiq-uitin-like modifier (SUMO) is actively involved in the pathogenesis of several diseases, including tumorigenesis, infections, immunity and neurodegenerative diseases, and has been considered a novel potential target for cancer treatment and other diseases. The mammalian cell genome contains five SUMO genes, SUMO1–5. SUMO4 and SUMO5 are restricted to specific tissues related to pathogenesis, such as diabetes3,4. SUMOs exist in the form of precursor protein and mature by cleaving the C-terminal amino acids by the specific protease (SENP) family (Fig 1. A)5,6. Mature SUMOs are conjugated to the target substrates through an enzymatic cascade reaction catalyzed by E1 activating enzyme, E2 conjugating enzyme and E3 ligase7-9. E1 activates mature SUMO through two main steps: first, Aos1 catalyzes adenylation of SUMO at C-terminus using ATP, and a covalent thioester bond is then formed between the and the C-terminal of SUMO and catalytic cysteine of Uba2. Subsequently, the activated SUMO is transferred to E2 (Ubc9) to form a thioester bond between the catalytic group cysteine of Ubc9 and SUMO10. Several E3s for SUMOylation (e.g., the PIAS family of proteins and RANBP2) are generally thought to determine substrate specificity in vivo and to conjugate SUMO to lysine residues of the substrates11,12. Conjugated SUMOs are then removed from substrates by the sen-trin-specific proteases (SENPs) to start the cycle again (Fig 1. A). In the SUMOylation cascade, many protein-protein interactions among SUMOs and its catalytic enzymes are critical in the successive conjugation and de-conjugation cascades, such as Aos1-Uba2, Uba2-Ubc9, and Ubc9-PIAS. We are interested in these interactions in both affinity and kinetics and would to understand the complicated conjugation processes so we can ma-nipulate this pathway to diagnose and treat diseases.

**Fig. 1.**
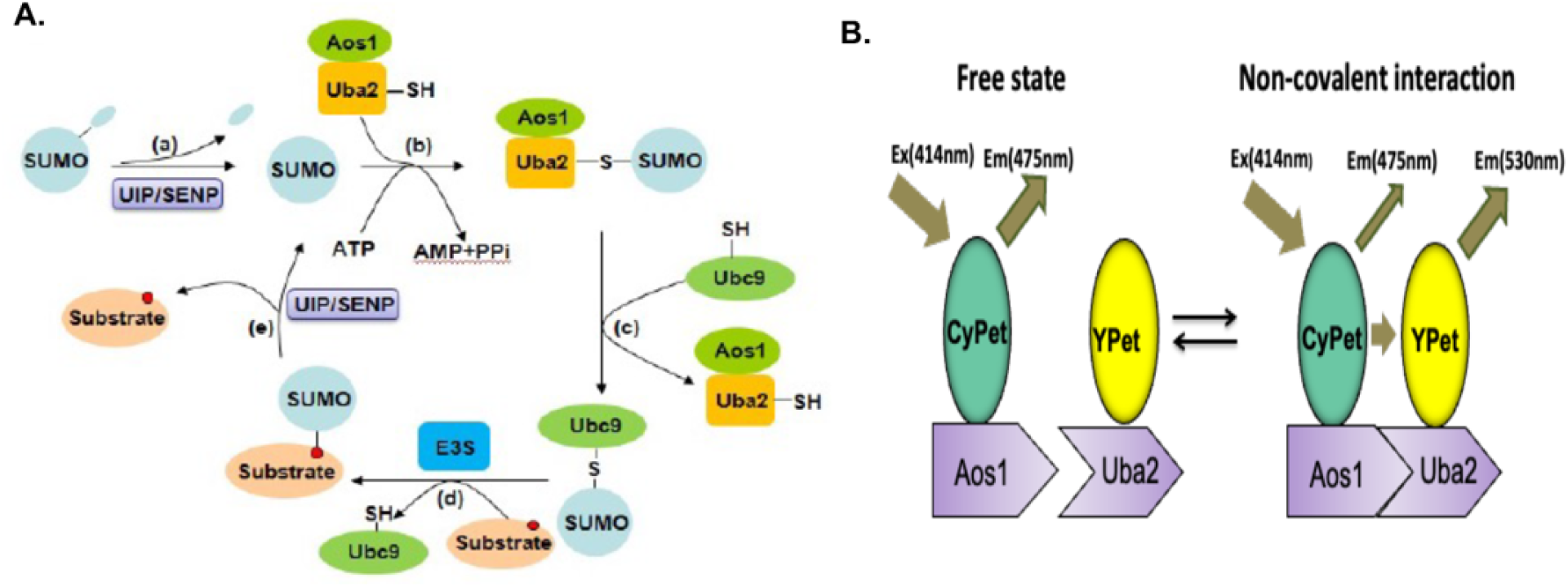
Schematic diagram of the SUMOylation cascade and FRET-based assay of the E1 heterodimer interaction. A. Diagram of SUMOylation cycle: (a) Cleavage of SUMO C-terminus; (b) SUMO activation and thioester bonding by E1 (Aos1-Uba2 dimmer); (c) SUMO transfer to Ubc9; (d) SUMOylation of substrate with or without the help by E3; (e) DeSUMoylation. B. Diagram of FRET within the SUMO E1 ligase resulting from Aos1 and Uba2 interaction.

Förster resonance energy transfer or fluorescence energy transfer (FRET), a mech-anism of energy transfer between two fluorescence molecules through nonradiative di-pole–dipole coupling, which is sensitive to the distance between a donor and acceptor within 1–10 nm. FRET has been widely used in bioengineering research for both optical imaging and spectroscope for analysis of protein interactions13-19. Because the FRET signal is proportional to the number of interaction events, considerable efforts have been made to develop it into a quantitative assay13, 20-22. However, due to the complexity of fluorescence emission from the donor and acceptor at the emission wavelength, these efforts have not been very successful. Our research team has successfully developed a quantitative FRET or qFRET method using a principle of cross-wavelength correlation co-efficiency approach to dissect the absolute FRET signal from noise to use this method for real-time protein-protein interaction signal change monitoring. Finally, after trying a variety of fluorescent pairs, a breakthrough has been made in the application of CyPet and YPet fluorescent pairs23,24. This method can determine absolute FRET signal and signals from free donor and acceptor at the emission wavelength in a single assay. A new mathematical formula for correlating the FRET signal with Kd was derived. In one of our previous studies of SUMO1-Ubc9 interactions, the Kd values of 0.26–0.31 μM for four different concentrations of CyPet-SUMO1 with YPet-Ubc9 were determined, in good agreement with those measured by the SPR method (0.35 μM) and isothermal titration calorimetry (ITC) (0.25 μM)25. The FRET pair CyPet/YPet we used also offered greater sensitivity26,27.

In a systems biology approach to understanding the multienzyme-catalyzed SUMO conjugation reaction, we are applying our novel qFRET assay to determine all the protein interaction affinities in the SUMOylation cascade. Here, we report the development of a qFRET assay for the determination of affinities between the E1 heterodimer subunits Aos1 and Uba2 with or without ATP and SUMO. The calculated Kd between Aos1 and Uba2 was in good agreement with that determined by SPR. These results not only provide new quantitative insight into the Aos1 and Uba2 interactions in the SUMOylation cascade, but also determine the interaction affinity changes of Aos1-Ub2 before and after SUMO1 activation for the first time.

## Materials and Methods

### DNA constructs, Protein expression, purification, and concentration measurement

The genes including Aos1, Uba2, SUMO1, CyPet, YPet, CyPet-Aos1, YPet-Uba2, and all vectors, pCRII-TOPO vector (Invitrogen), pET28(b) vector (Novagen), fused genes were provided by Jiayu Liao’s Lab. Protein expression, purification, and concentration measurement determination method can refer to the literature we published^28^.

### FRET measurements and analysis

The CyPet-Aos1 and YPet-Uba2 were mixed and incubated at 37°C in Tris buffer, including 20 mM Tris-HCl, 50 mM NaCl, adjusting pH to 7.5, in a total volume of 60 µL. The different concentration of CyPet-Aos1 was at 0.2, 0.5, 1, or 1.5 µM. The concentrations of YPet-Uba2 gradually increased from 0 to 5 µM. The reaction was stopped at 10 min, and the fluorescence multi-well plate reader FlexstationII384 (Molecular Devices, Sunnyvale, CA) was used for measurement. One of the fluorescence emission signals at 475 and 530 nm were collected when excited at 414 nm with a cutoff filter at 455 nm. Another fluorescence emission signal was collected at 530 nm when excited at 475 nm with a cutoff filter at 515 nm. Each data was repeated three times to take the average value and brought into the formula for calculation.

Before all the protein added to the plate, all the well should be scanned for blank. Then to determine the value of the ratio constants α and β to ascertain absolute E_mFRET_, when we determine the ratio constant α, a series of CyPet-Aos1 solutions was prepared at concentrations of 0.2, 0.5, 1.0, and 1.5 µM. Emission of CyPet-Aos1 at 475 and 530 nm was determined when excited at 414 nm. Dividing the emission at 475 nm (FL_DD_) by its emission at 530 nm obtained the ratio constant α. This is an estimate of the ratio of unquenched CyPet-Aos1 to total emission at 530 nm when excited at 414 nm. In the second series of experiments, YPet-Uba2 was prepared at 0.2, 0.5, 1.0 and 2.0, 3.0, 4.0, and 5.0 µM. Emission of YPet-Uba2 at 530 nm was determined when excited at 414 or 475 nm. Ratio constant β was obtained by dividing the YPet-Uba2 emission signal at 530nm when excited at 414 nm by the YPet-Uba2 signal at 530 nm when excited at 475 nm (FL_AA_).

### Data processing and K_d_ determination

After E_mFRET_ signals for all conditions were collected by FRET assay, the datasets of E_mFRET_(instrument collected) and the total concentrations of YPet-Uba2 (B-series)(designed) were fitted by Prism 5 to derive the value of E_mFRETmax_ and *K*_*d*_ according th following equation.

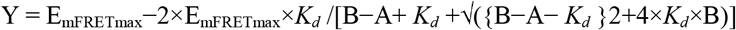

The parameters E_mFRET_, *K*_*d*_ and A are initial values equal to 1.0. Under the default constraints, E_mFRETmax_ must be greater than 0. A constant was used for concentration (0.2, 0.5, 1, and 1.5 µM). The mean ± standard deviation is the result.

### SPR determination of K_d_ for the non-covalent interaction of Aos1 with Uba2

All analyses of interactions between CyPet-Aos1 and YPet-Uba2 or Aos1 and Uba2 were performed on a BIAcore X100 system. NTA chip was used to detect the protein binding and dissociation experiment with his tag. Here we prepared three kinds buffer: Running buffer, including HEPES 10 mM, EDTA 50 μM, NaCl 150 mM, Tween20 0.005%, pH 7.4. Immobilizing buffer, including 500 µM NiCl2 in running buffer. Regeneration buffer, including HEPES 10 mM, EDTA 350 mM, NaCl 150 mM, Tween20 0.005%, pH 8.3. His-tagged CyPet-Aos1 and YPet-Uba2 or His-tagged Aos1 and Uba2 were dialyzed overnight in running buffer to be the same condition. The mobile phase rate is set to 30 µL/min. The experimental process is as follows: First, the NTA chip was treated with immobilizing buffer before immobilized 2 µg/mL purified CyPet-Aos1, or 1 µg/mL purified Aos1 protein. The CyPet-Aos1 or Aos1 was injected for 120 s and stabilized for 120 s. Then, YPet-Uba2 or Uba2 without His tag was injected by the rate of 30–120 µg/mL or 5–50 µg/mL respectively for 120 s and disassociated for 10 min. To continuously monitor nonspecific background binding of samples to the NTA surface, YPet-Uba2 and Uba2 proteins were injected into a control flow cell without treated by NiCl_2_ and CyPet-Aos1/Aos1 proteins. After monitoring one concentration of CyPet-Aos1 and YPet-Uba2 or Aos1 and Uba2, we regenerated the NTA sensor chip with regeneration buffer, then retreated with immobilizing buffer for another concentration. According to the requirements of the instrument, the experimental temperature is 25 °C. BIAcore X100 evaluation software ver. 1.0 was used to process data analysis.

### Aos1-Ubc2 interaction assay

For the assays of the interaction between Aos1 and Uba2 in the presence of SUMO1 and ATP, 0.6 μM CyPet-Aos1 and 0–5 μM YPet-Uba2, respectively were mixed in 40 μL buffer containing 50 mM Tris-HCl, 4 mM MgCl_2,_ and 1 mM DTT, pH 7.4. Fluorescence signals were monitored after 10 min. Then 10 μl of 1 μM SUMO1 was added to each well and fluorescence was monitored after 10 min., followed by 10 μl of 2 mM ATP added in the assay with monitoring of fluorescence at different time points. The samples were incubated at 37°C and measurements were made of E_mFRETmax_ and the FRET index.

## Results

### The Determination of E1 heterodimer K_d_, and the absolute FRET signal

The SUMO E1 heterodimer, Aos1 and Uba2, is a key enzyme in the SUMOylation pathway, and we therefore expanded our method of FRET-based *K*_*d*_ determination of CyPet-Aos1 and YPet-Uba2. We used FRET analysis to convert the concentration of binding protein and free protein into FRET signal intensity, and selected efficient FRET pairs CyPet and YPet^26,28,29^, which were fused with Aos1 and Uba2 respectively (Fig 1. B).

The general law of mass action for the interaction between Aos1 and Uba2 is as follows:

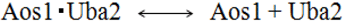

Since CyPet and YPet are fused with Aos1 and Uba2 respectively, the law can be written as follows:

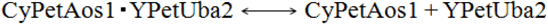

*and K*_*d*_ can be expressed as the following equation:

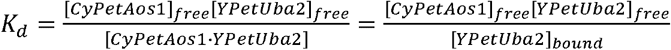

Since the FRET signal intensity changes with the concentration of CyPet-Aos1 and YPet-Uba2, the left and right sides of the following equation are equal:

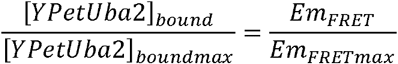

According to the method of FRET established by our experimental team^23,24^, *K*_*d*_ can be finally obtained through the following 10 equations:

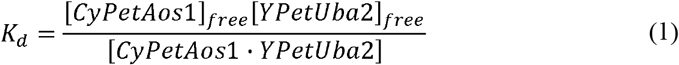

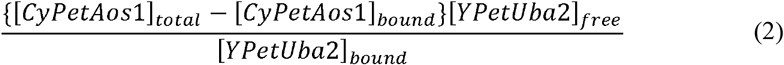

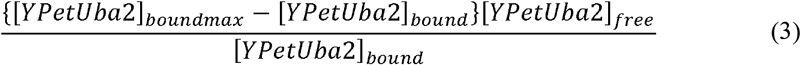

Equation (3) can be converted to:

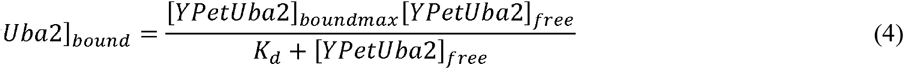

It can be deduced from equations (2) and (4):

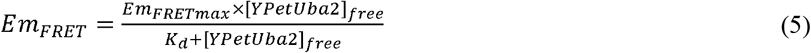

Here, A is defined as the total concentration of CyPet-Aos1 ([CyPet-Aos1]_total_), B is the total concentration of YPet-Uba2 ([YPet-Uba2]_total_), and Y is the concentration of free YPet-Uba2 ([YPet-Uba2]_free_). We can write [YPet-Uba2]_bound_ and [CyPet-Aos1]_free_ as the following formula:

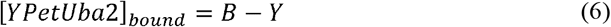

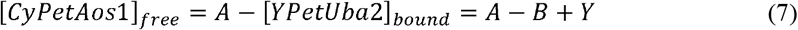

According to the equations (6) and (7), *K*_*d*_ can be deduced that:

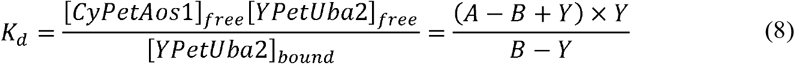

Then the equation can be solved to obtain:

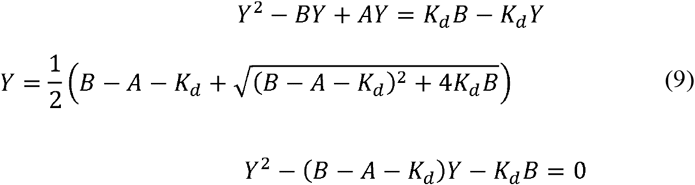

Combining equations (5) and (9), it can be concluded that:

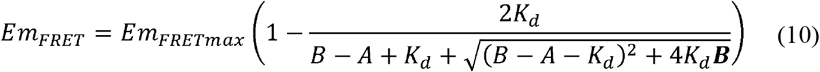

Here, A and B are the known concentrations set in the experiment. E_mFRET_ can be obtained from the FRET results measured by the instrument, and the unknown parameters are E_mFRETmax_ and *K*_*d*_.

Absolute FRET signals of CyPet-Aos1 and YPet-Uba2 is measured by the method established in our laboratory^23,24^.

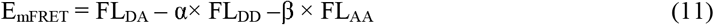

In the FRET analysis for determining the interaction between CyPet-Aos1 and YPet-Uba2, the ratio constants α is the ratio of the emission signal value of CyPet-Aos1 at 530 nm to the fluorescence signal intensity at 475 nm when excited at 414 nm, and β is the ratio of the fluorescence signal intensity of YPet-Uba2 at 530 nm when excited at 414 nm to the 530 nm emission wavelength when excited at 475 nm. Here *α* and *β* in the equation are determined to be 0.334±0.003 and 0.026±0.004, respectively (Fig 2).

**Fig 2.**
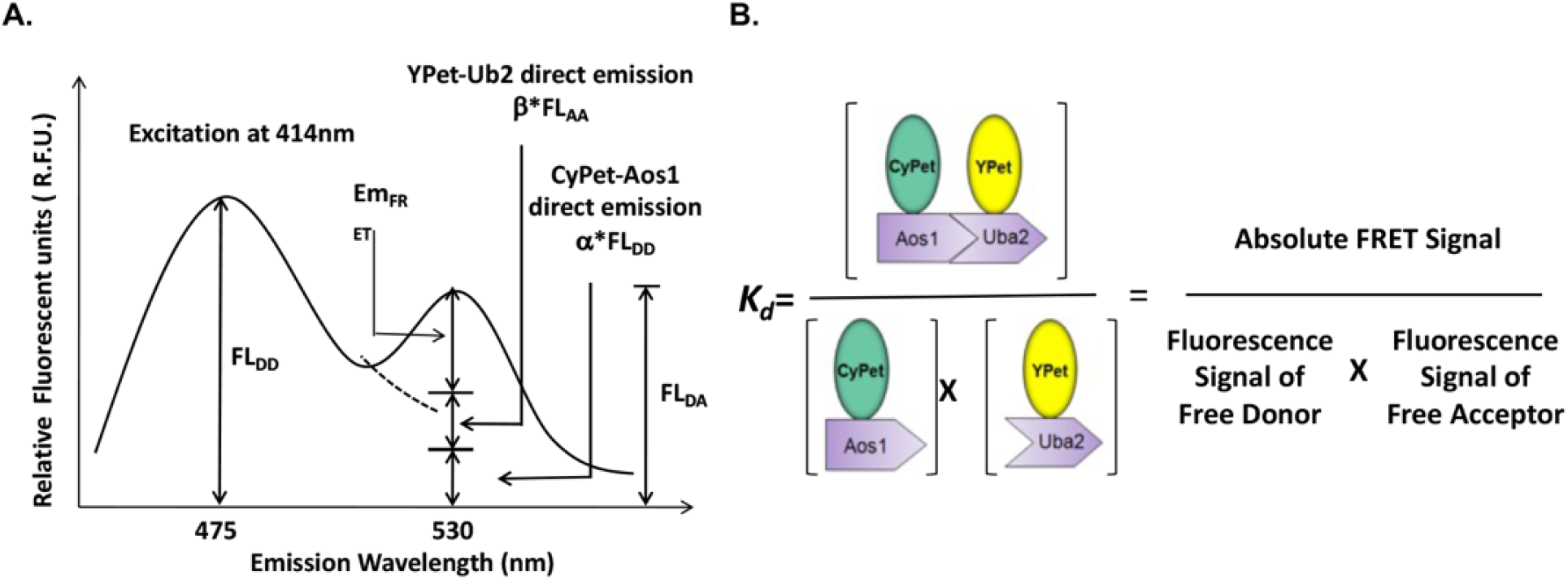
Differential analysis of fluorescence signals in a FRET assay and FRET-based *K*_*d*_ determination. A. Fluorescence emission at 530 nm (FL_DA_) is divided into three parts: FRET from acceptor YPet-Uba2(E_mFRET_), direct emission from the donor CyPet-Aos1 (α×FL_DD_), and direct emission from the acceptor YPet-Uba2 (β×FL_AA_) ^23,24^. B. *K*_*d*_ is represented by the fluorescence signals from the bound pair divided by the product of the two fluorescence signals from the free proteins.

The sensitivity of FRET analysis under different concentrations of CyPet-Aos1 was detected. In one set of experiments, the CyPet-Aos1 concentration was fixed at 1 µM (Fig 3. A). When the concentration of YPet-Uba2 was 0, the emission peak of CyPet-Aos1 appeared at 475 nm and the fluorescence value was low at 530 nm. As the concentration of YPet-Uba2 gradually increased from 0 to 5 µM, FRET occurred due to the proximity between YPet-Uba2 and CyPet-Aos1, resulting in a significant increase in the emission fluorescence at 530 nm and a gradual decrease in the fluorescence intensity at 475 nm. To verify whether the *K*_*d*_ data obtained by equations (10) and (11) is stable. We determined FRET signal changes of four different concentrations of CyPet-Aos1, 0.2, 0.5, 1.0, 1.5 µM respectively, and the concentration of YPet-Uba2 increased from 0 µM to 5 µM. The result showed that as more YPet-Uba2 was added to the assay, E_mFRET_ gradually increased to E_mFRETmax_ (Fig 3. B(a-d)).

**Fig 3.**
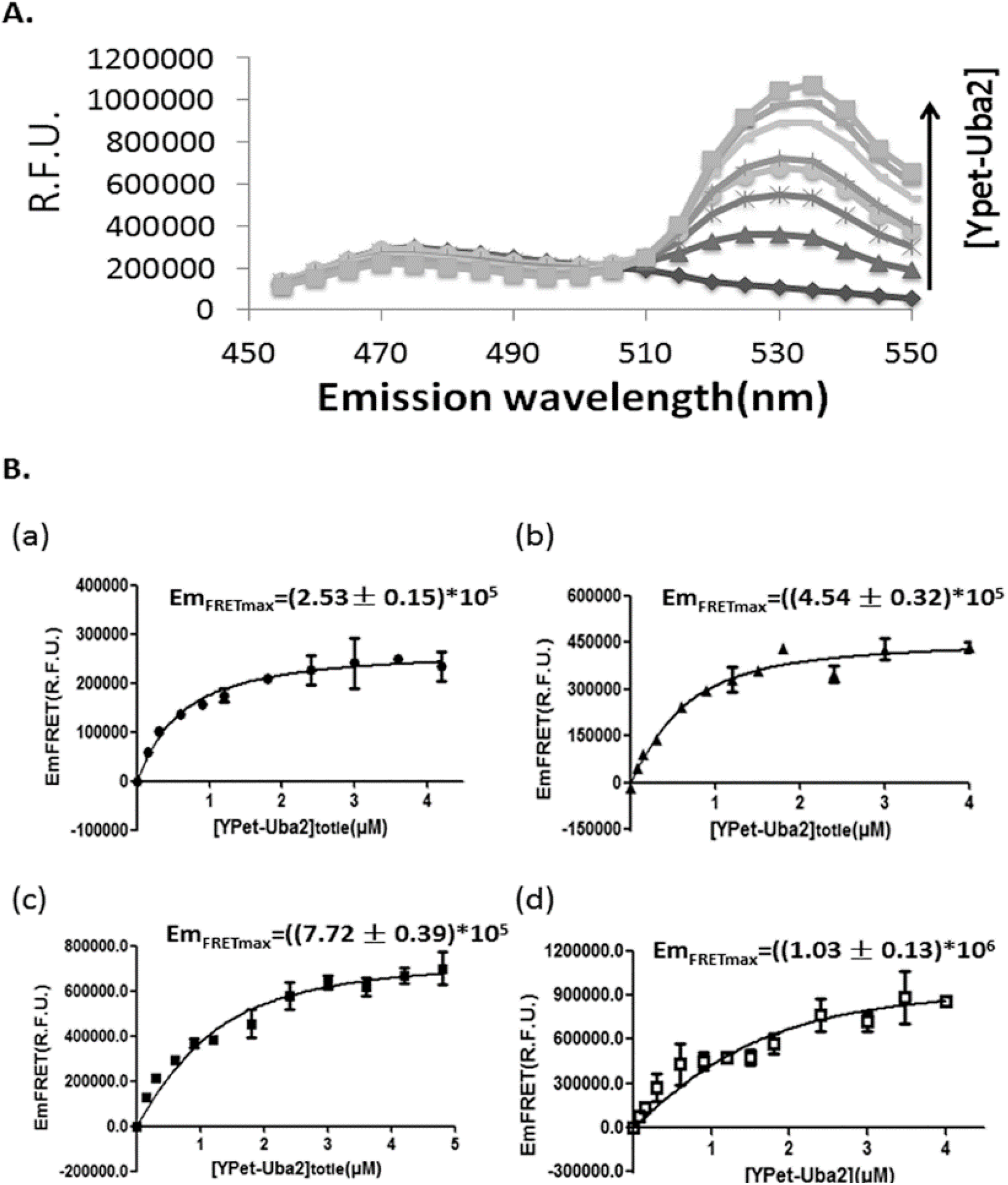
FRET assay for Aos-Uba2 interaction. A. The FRET intensity titration with increasing concentrations of YPet-Uba2. B.(a)-(d). E_mFRET_ and E_mFRETmax_ determinations at different concentrations of CyPet-Aos1. Plots of E_mFRET_ and E_mFRETmax_ determinations at (a) 0.2, (b) 0.5, (c) 1.0 and (d) 1.5 µM of CyPet-Aos1 with increasing concentration of YPet-Uba2.

### Determination of the E1 interaction affinity, K_d_, by FRET assay

After the concentration of CyPet-Aos1, YPet-Uba2 and Em_FRET_ data, brought into equation(10), *K*_*d*_ and E_mFRETmax_ can be obtained through prism 5 software. Therefore, through nonlinear regression, the measured values of E_mFRETmax_, were 2.53 (± 0.15) × 10^5^, 4.54 (± 0.15) × 10^5^, 7.72 (± 0.39) × 10^5^, and 1.03 (± 0.13) × 10^6^ RFU. E_mFRETmax_ was linearly correlated with the concentration of CyPet-Aos1 in our assay, and E_mFRETmax_ (Table 1) is plotted against the corresponding concentration of CyPet-Aos1 in Fig 4. A and B (R^2^= 0.997).

**Table 1.** Summary of maximal FRET emission E_mFRETmax_ and *K*_*d*_ values

| [CyPet-Aos1]<sub>total</sub> (μM) | 0.2 | 0.5 | 1.0 | 1.5 |
| --- | --- | --- | --- | --- |
| E<sub>mFRETmax</sub> (RFU) | (2.53 ± 0.15)×10<sup>5</sup> | (4.54 ± 0.15)×10<sup>5</sup> | (7.72 ± 0.39)×10<sup>5</sup> | (1.03 ± 0.13)×10<sup>6</sup> |
| K<sub>d</sub> (μM) | 0.48±0.12 | 0.31±0.10 | 0.55±0.13 | 0.55±0.32 |

**Fig 4.**
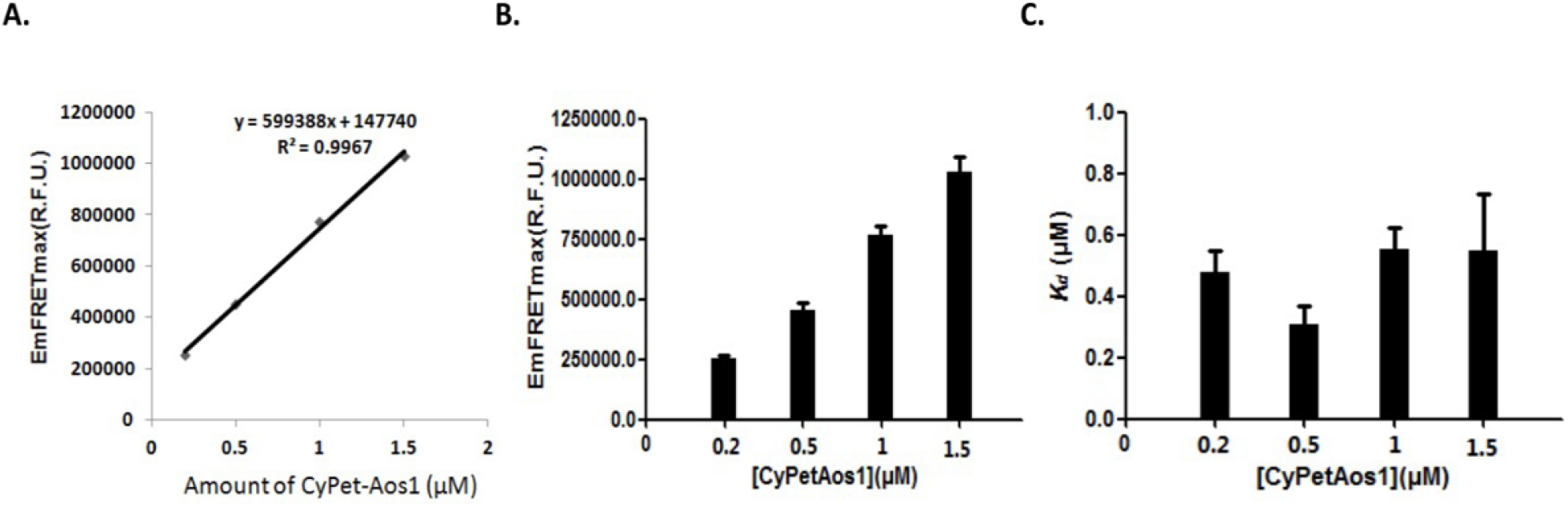
Determinations of E_mFRETmax_ and *K*_*d*_ at different concentrations of CyPet-Aos1. A. Maximal FRET strength is directly proportional to the concentration of CyPet-Aos1. B. Column chart of E_mFRETmax_ vs. concentration of CyPet-Aos1. C. *K*_*d*_ values at different amount of CyPet-Aos1.

The dissociation constant between CyPet-Aos1 and YPet-Uba2, *K*_*d*_, were 0.48±0.12 μM, 0.31±0.10 μM, 0.55±0.13 μM, and 0.55±0.32 μM, respectively (Fig 4. C and Table 1) under the four concentrations of CyPet-Aos1 (0.2, 0.5, 1.0, and 1.5 μM). These results suggest that we can calculate *K*_*d*_ using the qFRETassay, and the values of *K*_*d*_ determined by this method are consistent and accurate. The very similar values of *K*_*d*_ calculated from the different concentrations of CyPet-Aos1 and YPet-Uba2 (0.2 to 1.5 μM donor, and at concentrations of 3–20 times its binding acceptor) also indicate that the FRET-based approach to *K*_*d*_ measurement is very stable and reliable.

### Determination of the E1 interaction affinity, K_d_, by SPR

To validate our results, we processed the assay for dissociation constant between CyPet-Aos1 and YPet-Uba2 by SPR. CyPet-Aos1 was tagged by His. NTA sensor chip was used to immobilized his-tagged CyPet-Aos1. When different concentrations of YPet-Uba2, 30, 60, 90, 90, 120 μM, pass through the chip with the mobile phase, it interacts with CyPet-Aos1 and causes the change of response signal (Fig.5A). The *K*_*d*_ value determined by SPR method is 0.93 µ M. This data is very close to that determined by FRET method. The *K*_*d*_ of Aos1 and Uba2 without labeled fluorescence pair CyPet/YPet was also determined by SPR method (Fig.5B), and the result was 0.96 μM. Various verification methods shows that the determination of *K*_*d*_ by FRET method is very accurate, but the cost is much lower than SPR method, and is much faster. Furthermore, the FRET method can be used to determine the optimum temperature of the enzyme at 37°C, which is the temperature that Biacore instrument cannot set.

**Fig 5.**
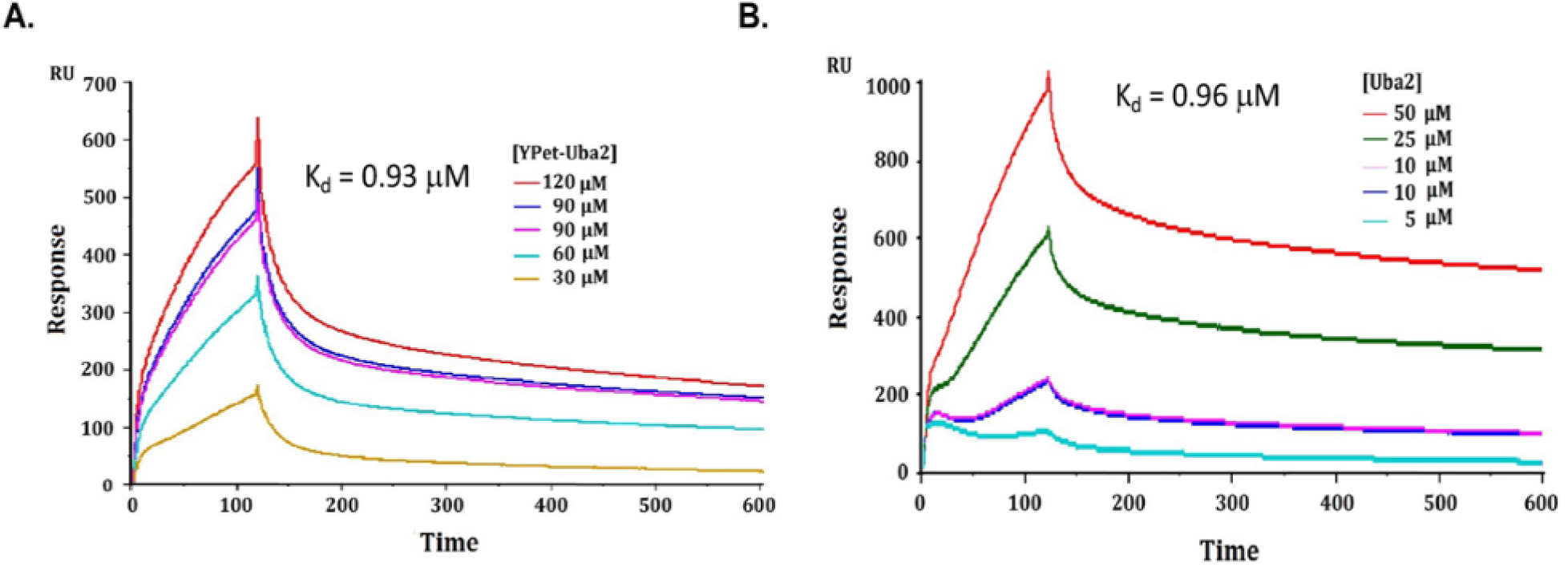
Determination of interaction affinity *K*_*d*_ by SPR. A. *K*_*d*_ between CyPet-Aos1 and YPet-Uba2 interaction, 0.93 µM. B. *K*_*d*_ between Aos1 and Uba2, 0.96 µM.

#### Aos1-Uba2 interaction in the activation cascade

In the SUMO pathway, E1 transfers sumo to Uba2, but it is not clear how the equilibrium dissociation of E1 (Aos1/Uba2) changes during this process. This interaction can be monitored in real-time by FRET when ATP is added to SUMO. Seven regression plots corresponding to one concentration of CyPet-Aos1(0.6 μM) over the range of YPet-Uba2 concentrations (0-5 μM) used in our experiments are shown in Fig.6. From the non-linear regression calculated in Prism 5, the predicted values of E_mFRETmax_ were 5.86 (± 0.40) × 10^5^, 7.59 (± 0.42) × 10^5^, 4.71 (± 0.18) × 10^5^, 4.77 (± 0.17) × 10^5^, 4.86 (± 0.15) × 10^5^, 5.47 (± 0.18) × 10^5^ and 5.82 (± 0.14) × 10^5^. The values of *K*_*d*_ calculated from these seven conditions were 0.4261±0.1531, 0.8422±0.1684, 0.2189±0.0641, 0.2144±0.0619, 0.1194±0.0425, 0.1661±0.0505, 0.1104±0.0324 (Fig.7, Table 2). Under different conditions, after SUMO1 and ATP were added to the assay, E_mFRET_ and the FRET index changed (Fig.8). The data showed that when SUMO1 was added into the system, *K*_*d*_ increased. It can be inferred from this result that SUMO1 interacts with Aos1 or Uba2 during this process, leading to dimmer separation. When ATP was added after SUMO1, *K*_*d*_ decreased. It can be inferred that Aos1 and Uba2 quickly formed a dimmer, which catalyzed the formation of a thioester bond between Uba2 and SUMO1. The *K*_*d*_ data show that the equilibrium between Aos1 and Uba2 favors dimmer formation, and therefore the E1 dimmer is more stable.

**Table 2.** Summary of *K*_*d*_ values and maximal FRET emission E_mFRETmax_ with or without ATP and SUMO1

| Different assay | $K_d$ (μM) + SD | $E_{mFRETmax}$ (R.F.U.) + SD |
| --- | --- | --- |
| a: Aos1+Uba2 | 0.43±0.15 | 585933±40098 |
| b: Aos1+Uba2+SUMO1 | 0.84±0.17 | 758647±42057 |
| c: Aos1+Uba2+SUMO1+ATP 5min | 0.22±0.06 | 470567±17724 |
| d: Aos1+Uba2+SUMO1+ATP 10min | 0.21±0.06 | 477464±17489 |
| e: Aos1+Uba2+SUMO1+ATP 20min | 0.12±0.04 | 485560±15288 |
| f: Aos1+Uba2+SUMO1+ATP 30min | 0.17±0.05 | 547385±18052 |
| g: Aos1+Uba2+SUMO1+ATP 60min | 0.11±0.03 | 582227±14406 |

**Fig 6.**
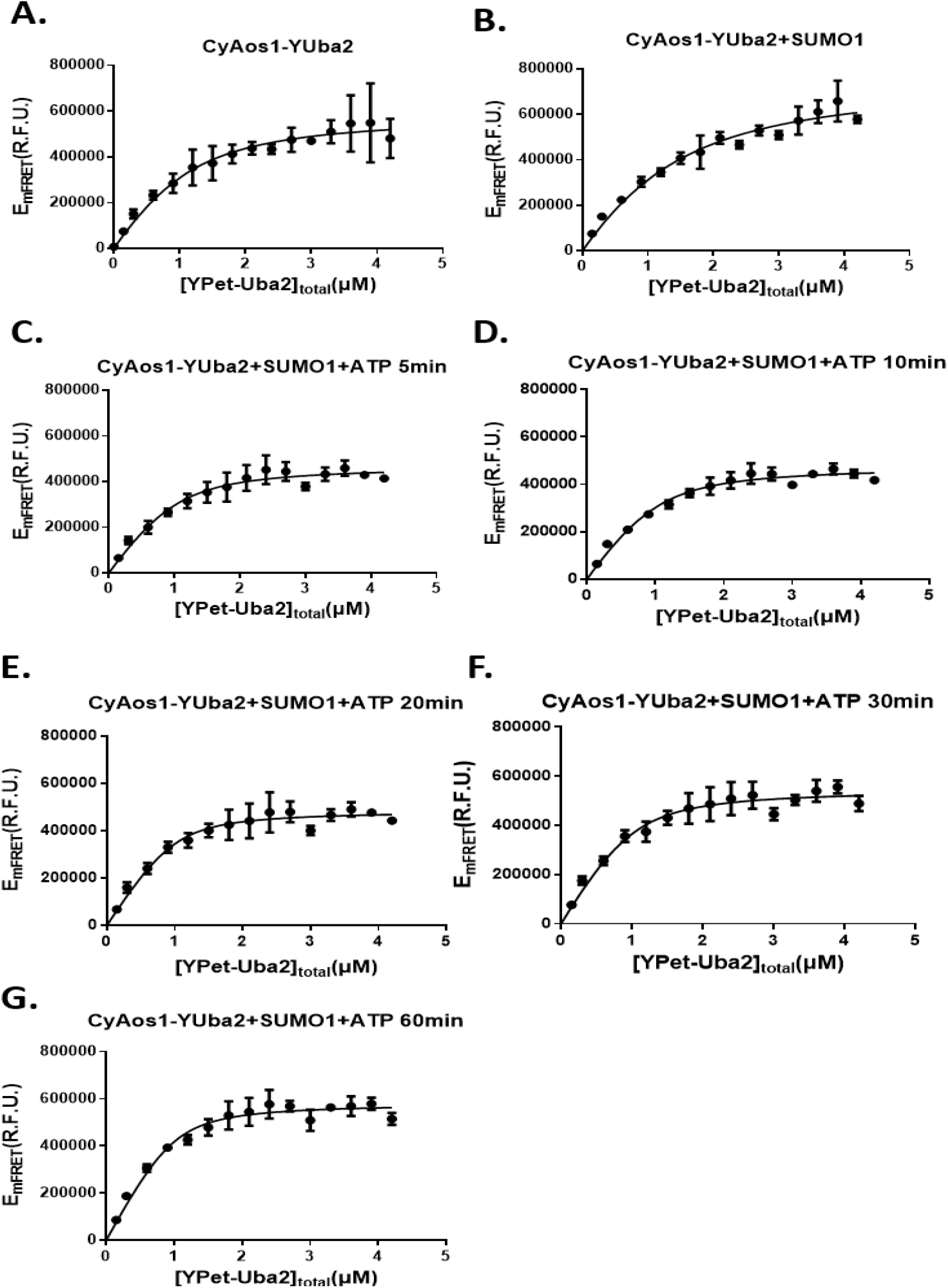
Plot of E_mFRET_ vs. [YPet-Uba2] in the presence of SUMO1 and ATP.

**Fig 7.**
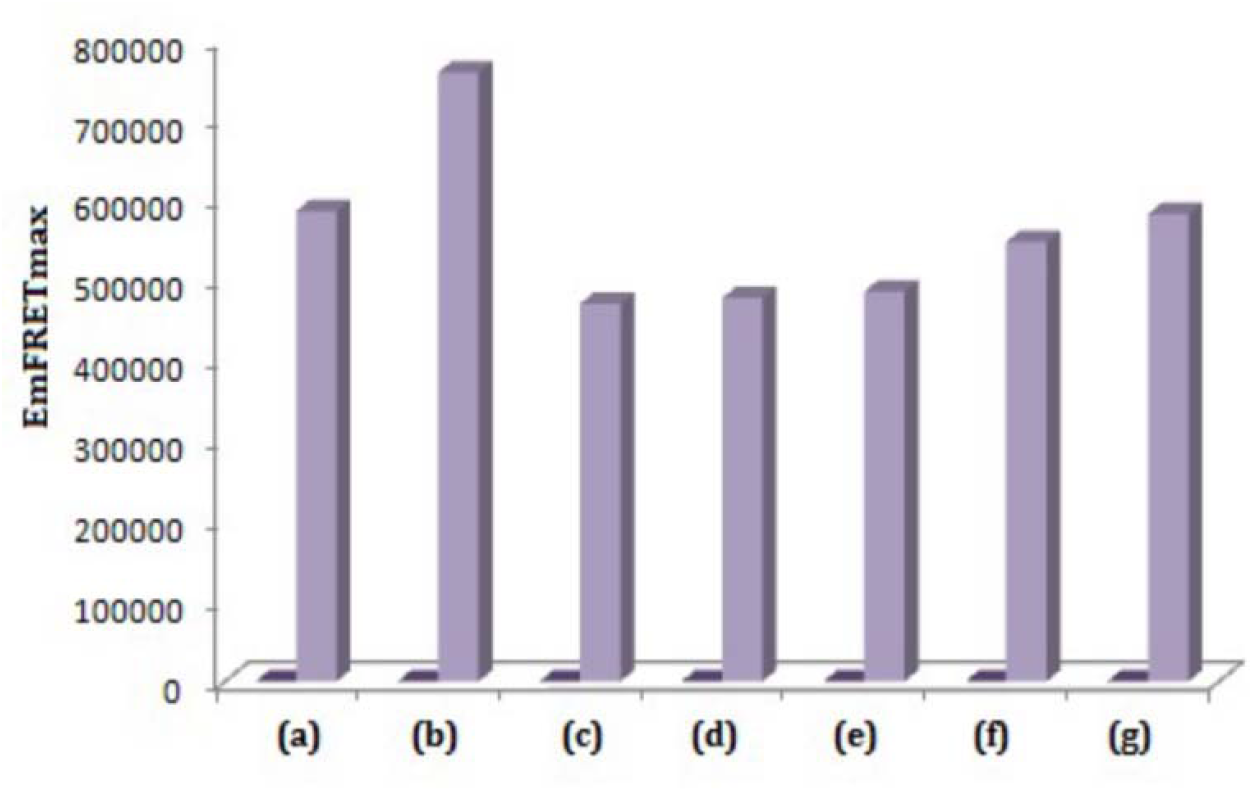
Plot E_mFRET_ for different reaction systems. (a) CyPet-Aos1-YPet-Uba2, 10 min at 37°C; (b) CyPet-Aos1-YPet-Uba2+SUMO1, 10 min at 37°C; (c) CyPet-Aos1-YPet-Uba2+SUMO1+ATP, 5 min at 37°C; (d) CyPet-Aos1-YPet-Uba2+SUMO1+ATP, 10 min at 37°C; (e) CyPet-Aos1-YPet-Uba2+SUMO1+ATP, 20 min at 37°C; (f) CyPet-Aos1-YPet-Uba2+SUMO1+ATP, 30 min at 37°C; (g) CyPet-Aos1-YPet-Uba2+SUMO1+ATP, 60 min at 37°C.

**Fig 8.**
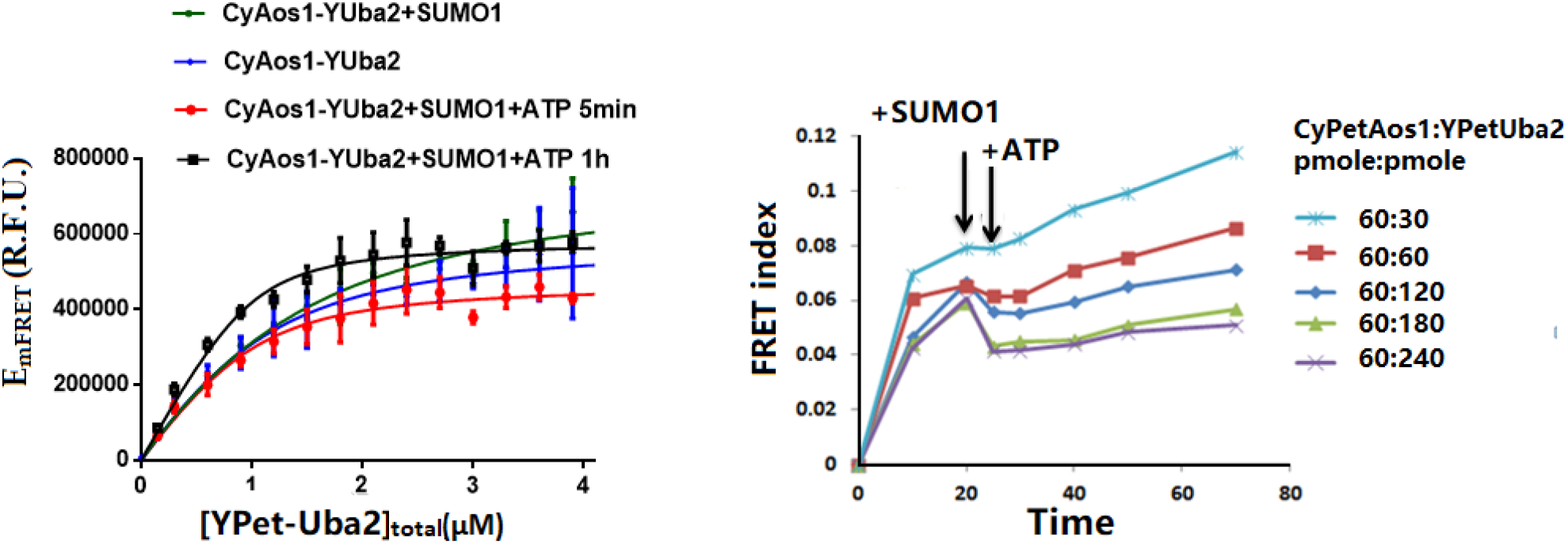
**A**. Variation of E_mFRET_ after adding SUMO1 and ATP in the assay. B. Plot of FRET index vs. time.

## Discussion

Here we report the use of a qFRET assay to determine the dissociation constant, *K*_*d*_, between Aos1 and Uba2 in the absence and presence of SUMO1 and ATP for the first time. This finding provides valuable information on the interaction affinity between the UBL E1 heterodimer subunits for the first time. The *K*_*d*_ values ranging from 0.31 to 0.55 μM, determined using concentrations of CyPet-Aos1 ranging from 0.2 to 1.5 μM, are very consistent with those determined by traditional SPR (0.93 for CyPet-Aos1 and YPet-Uba2, and 0.96 μM Aos1 and Uba2). The slight difference in *K*_*d*_ values between these two approaches may be due to three reasons. First, the *K*_*d*_ determination using qFRET was conducted in solution, while the K_d_ determination using SPR was conducted at the surface of the glass, which may affect protein conformation; Second, there were some impurities in Uba2 that can affect SPR assay results; Third, the buffer and the temperature used for the two methods are different. However, in our qFRET assay, we determined the protein concentration by the Bradford method and fluorescence measurements of both the fusion proteins and the fluorescent proteins alone. The temperature used in our method is 37°C, which is the optimum temperature for an enzyme reaction. The detection process is very fast, which greatly reduces the risk of denaturation of enzymes in the environment. Meanwhile, the real-time quantitative monitoring of Aos1-Uba2 using qFRET in solution provides more physiological conditions for understanding the multistep cascade reaction of SUMO modification.

In the presence of SUMO1 and ATP, the change in *K*_*d*_ can also be determined in real-time. The interaction affinity of the SUMO1 E1 heterodimer before and after activation can provide more kinetics parameters to understand better the chemical mechanisms of adenylation and thioester bond formation. Although the E1 heterodimer co-crystallization with the SUMO peptide provided some insights into the complex interactions and conformational changes during the first step of activation, the affinity changes of the E1 heterodimer interaction during the activation step was not previously determined. The affinity changes of Aos1/Uba2 are very challenging to determine using other methods. The observed moderate interaction affinity before SUMO1 activation is consistent with general protein interactions in cells and may balance E1 heterodimer interactions and subsequent conformational changes to increase interaction affinity during adenylation, followed by thioester bond formation. A previous study of SUMOylation E1 enzyme kinetics suggests that high-affinity interactions may not favor subsequent conformational changes^30^. This also holds true for other UB/UBL E1 family members as well as other enzymatic reactions. It will be exciting to determine whether the SUMO E1 heterodimer interaction affinity change favors adenylation and thioester bond formation. This information should provide insights into the mechanisms of the complex SUMO activation cascade and will also provide opportunities for future therapeutic developments targeting SUMOylation and other UBLs.

In conclusion, in the study, we carried out the detection of Aos1/Uba2 protein-protein interaction based on receptor emission FRET assay, by which is numerically close to SPR method. However, it gets more advantages: First, it is an accurate and sensitive approach. Second, it is the simplicity of experimental operation. Third, it is easy to make accurate temperature control. Forth, the protein dosage is very low to picomole, and experimental cost is lower^29^. In addition, by adding other substances (SUMO1 and ATP in this experiment) in detecting protein interaction, the influence of these substances on the equilibrium dissociation of protein interaction can be detected in real time. It is hoped that this technology can help more researchers complete their work in the protein-protein interaction field in the future.

## Author Contributions

Conceptualization, J.L. and L. J.; methodology, J.L. and L. J; Validation, L. J; formal analysis, J.L. and L. J; investigation, J.L. and L. J; resources, J.L. and L. J; data curation, L.J. writing—original draft preparation, L.J.; writing—review and editing, Y.T., X.W. and J.L.; supervision, J.L.; project administration, J.L.; funding acquisition, J.L. and L. J. All authors have read and agreed to the published version of the manuscript.

## Data Availability Statement

The data supporting this article have been included as part of the Supplementary Information.

## Acknowledgments

We are very grateful to Dr. Songqin Pan in the Institute for Integrative Genome Biology for very valuable helps in Biacore instrument and trouble-shootings. We thank all the members in Liao’s group for very close collaborative work and helps for the work.

## References

1 Abrieu, A. & Liakopoulos, D. How Does SUMO Participate in Spindle Organization? Cells 8, doi:10.3390/cells8080801 (2019).

2 Psakhye, I. & Jentsch, S. Protein group modification and synergy in the SUMO pathway as exemplified in DNA repair. Cell 151, 807–820, doi:10.1016/j.cell.2012.10.021 (2012).

3 Guo, D. et al. A functional variant of SUMO4, a new I kappa B alpha modifier, is associated with type 1 diabetes. Nat Genet 36, 837–841, doi:10.1038/ng1391 (2004).

4 Liang, Y. C. et al. SUMO5, a Novel Poly-SUMO Isoform, Regulates PML Nuclear Bodies. Sci Rep 6, 26509, doi:10.1038/srep26509 (2016).

5 Mukhopadhyay, D. & Riezman, H. Proteasome-independent functions of ubiquitin in endocytosis and signaling. Science 315, 201–205, doi:10.1126/science.1127085 (2007).

6 Yeh, E. T. SUMOylation and De-SUMOylation: wrestling with life’s processes. J Biol Chem 284, 8223–8227, doi:10.1074/jbc.R800050200 (2009).

7 Melchior, F. SUMO--nonclassical ubiquitin. Annu Rev Cell Dev Biol 16, 591–626, doi:10.1146/annurev.cellbio.16.1.591 (2000).

8 Johnson, E. S. Protein modification by SUMO. Annu Rev Biochem 73, 355–382, doi:10.1146/annurev.biochem.73.011303.074118 (2004).

9 Geiss-Friedlander, R. & Melchior, F. Concepts in sumoylation: a decade on. Nat Rev Mol Cell Biol 8, 947–956, doi:10.1038/nrm2293 (2007).

10 Johnson, E. S. & Blobel, G. Ubc9p is the conjugating enzyme for the ubiquitin-like protein Smt3p. J Biol Chem 272, 26799–26802, doi:10.1074/jbc.272.43.26799 (1997).

11 Kahyo, T., Nishida, T. & Yasuda, H. Involvement of PIAS1 in the sumoylation of tumor suppressor p53. Mol Cell 8, 713–718, doi:10.1016/s1097-2765(01)00349-5 (2001).

12 Gareau, J. R. & Lima, C. D. The SUMO pathway: emerging mechanisms that shape specificity, conjugation and recognition. Nat Rev Mol Cell Biol 11, 861–871, doi:10.1038/nrm3011 (2010).

13 Gordon, G. W., Berry, G., Liang, X. H., Levine, B. & Herman, B. Quantitative fluorescence resonance energy transfer measurements using fluorescence microscopy. Biophys J 74, 2702–2713, doi:10.1016/S0006-3495(98)77976-7 (1998).

14 Jones, J., Heim, R., Hare, E., Stack, J. & Pollok, B. A. Development and application of a GFP-FRET intracellular caspase assay for drug screening. J Biomol Screen 5, 307–318, doi:10.1177/108705710000500502 (2000).

15 He, L. et al. Monitoring caspase activity in living cells using fluorescent proteins and flow cytometry. Am J Pathol 164, 1901–1913, doi:10.1016/S0002-9440(10)63751-0 (2004).

16 Nguyen, A. W. & Daugherty, P. S. Evolutionary optimization of fluorescent proteins for intracellular FRET. Nat Biotechnol 23, 355–360, doi:10.1038/nbt1066 (2005).

17 Wu, X. et al. Measurement of two caspase activities simultaneously in living cells by a novel dual FRET fluorescent indicator probe. Cytometry A 69, 477–486, doi:10.1002/cyto.a.20300 (2006).

18 Zhang, L., Lei, J., Liu, J., Ma, F. & Ju, H. Persistent luminescence nanoprobe for biosensing and lifetime imaging of cell apoptosis via time-resolved fluorescence resonance energy transfer. Biomaterials 67, 323–334, doi:10.1016/j.biomaterials.2015.07.037 (2015).

19 Savitsky, A. P. et al. FLIM-FRET Imaging of Caspase-3 Activity in Live Cells Using Pair of Red Fluorescent Proteins. Theranostics 2, 215–226, doi:10.7150/thno.3885 (2012).

20 Saucerman, J. J. et al. Systems analysis of PKA-mediated phosphorylation gradients in live cardiac myocytes. Proc Natl Acad Sci U S A 103, 12923–12928, doi:10.1073/pnas.0600137103 (2006).

21 Dams, G. et al. A time-resolved fluorescence assay to identify small-molecule inhibitors of HIV-1 fusion. J Biomol Screen 12, 865–874, doi:10.1177/1087057107304645 (2007).

22 Chen, H., Puhl, H. L., 3rd & Ikeda, S. R. Estimating protein-protein interaction affinity in living cells using quantitative Forster resonance energy transfer measurements. J Biomed Opt 12, 054011, doi:10.1117/1.2799171 (2007).

23 Song, Y., Madahar, V. & Liao, J. Development of FRET assay into quantitative and high-throughput screening technology platforms for protein-protein interactions. Ann Biomed Eng 39, 1224–1234, doi:10.1007/s10439-010-0225-x (2011).

24 Song, Y., Rodgers, V. G., Schultz, J. S. & Liao, J. Protein interaction affinity determination by quantitative FRET technology. Biotechnol Bioeng 109, 2875–2883, doi:10.1002/bit.24564 (2012).

25 Tatham, M. H. et al. Role of an N-terminal site of Ubc9 in SUMO-1, -2, and -3 binding and conjugation. Biochemistry 42, 9959–9969, doi:10.1021/bi0345283 (2003).

26 Jiang, L. et al. Specific substrate recognition and thioester intermediate determinations in ubiquitin and SUMO conjugation cascades revealed by a high-sensitive FRET assay. Mol Biosyst 10, 778–786, doi:10.1039/c3mb70155g (2014).

27 Jiang, L. et al. Protein-Protein Affinity Determination by Quantitative FRET Quenching. Sci Rep 9, 2050, doi:10.1038/s41598-018-35535-9 (2019).

28 Liu, Y. et al. Product inhibition kinetics determinations - Substrate interaction affinity and enzymatic kinetics using one quantitative FRET assay. Int J Biol Macromol 193, 1481–1487, doi:10.1016/j.ijbiomac.2021.10.211 (2021).

29 Liao, J., Madahar, V., Dang, R. & Jiang, L. Quantitative FRET (qFRET) Technology for the Determination of Protein-Protein Interaction Affinity in Solution. Molecules 26, doi:10.3390/molecules26216339 (2021).

30 Olsen, S. K., Capili, A. D., Lu, X., Tan, D. S. & Lima, C. D. Active site remodelling accompanies thioester bond formation in the SUMO E1. Nature 463, 906–912, doi:10.1038/nature08765 (2010).

